# Characterization of the N7 RNA cap methyltransferase from *Trichomonas vaginalis* and inhibitor discovery

**DOI:** 10.64898/2026.09.09.750033

**Authors:** Jakub Benysek, Dominika Chalupska, Milan Stefek, Olga Bobileva, Tomas Otava, Milan Dejmek, Bartosz Różycki, Radim Nencka, Evzen Boura

## Abstract

RNA cap formation is essential for eukaryotic gene expression, yet the enzymes responsible for cap methylation remain poorly characterized in many eukaryotic parasites, including *Trichomonas vaginalis*. Here, we functionally and structurally characterized the RNA guanine-N7 methyltransferases (MTases) from *T. vaginalis, Tv*RNMT1 and *Tv*RNMT2. We determined the crystal structure of *Tv*RNMT1 in complex with S-adenosylhomocysteine (SAH) at 1.6 Å resolution and additional structures with the inhibitors sinefungin and OBO101, defining the architecture and ligand-recognition properties of its active site. Screening of SAH analogues identified several inhibitors with submicromolar to low-micromolar activity against both homologues. Molecular docking and modelling of capped RNA binding further delineated the inhibitor- and RNA-binding regions of *Tv*RNMT1 and identified residues that may contribute to cap recognition. These results define the molecular basis of ligand recognition by *T. vaginalis* RNA cap MTases and establish a structural framework for targeting parasite RNA capping.

## Introduction

*Trichomonas vaginalis* (*T. vaginalis*) is a pear-shaped protozoan parasite that causes trichomoniasis, the most prevalent non-viral sexually transmitted infection (STI). Approximately 156 million new cases were estimated globally in 2020 [1-4]. Motile T. vaginalis trophozoites carry four free anterior flagella and a recurrent fifth flagellum associated with the undulating membrane, which is essential for its movement. In women, *T. vaginalis* infection can lead to vaginitis, while in men, it may result in urethritis and prostatitis. Importantly, this infection increases the risk of HIV transmission and other sexually transmitted diseases (STDs) in both sexes. The current standard treatment involves 5-nitroimidazoles [5, 6]; however, the rise of drug-resistant strains presents a major public health concern due to the lack of alternative treatment options [7, 8]. As a result, there is an urgent need for effective antiparasitic compounds.

RNA capping is a fundamental and essential process in all eukaryotic organisms. The addition of a 5’ cap structure to nascent mRNA transcripts plays a critical role in RNA stability, efficient splicing, nuclear export, and translation initiation [9, 10]. Without proper capping, mRNAs are rapidly degraded by exonucleases, and their ability to be translated into functional proteins is significantly impaired. Consequently, RNA capping is essential for maintaining normal cellular function and overall organismal viability [9, 11].

The formation of the canonical cap 0 structure involves three enzymatic reactions. First, an RNA triphosphatase removes the γ-phosphate from the 5’ end of the nascent mRNA, leaving a diphosphate structure. Next, a guanylyltransferase catalyzes the attachment of a guanosine monophosphate (GMP) from GTP through an unusual 5’-to-5’ triphosphate linkage, forming the pre-cap structure. Finally, this structure is methylated at the N7 position of the guanine to generate cap 0. In most eukaryotes but, interestingly not in some lower eukaryotic organisms such as *S. cerevisiae*, the ribose 2′-O position is also methylated by a cap-specific 2′-O MTase, generating the cap 1 structure. The first two activities are encoded by separate proteins in many microbial eukaryotes, whereas in metazoans (including humans), plants and *T. vaginalis*, they reside in a single bifunctional protein [12], while the methylation steps are catalysed by two methyltransferases (MTases), RNA guanine-7 methyltransferase (RNMT) and cap methyltransferase (CMTR), respectively. Both of these enzymes, similarly to most MTases, use

S-adenosylmethionine (SAM) as the methyl donor. SAM is converted to S-adenosylhomocysteine (SAH) during the reaction and the methylated RNA is formed.

RNA capping is relatively well understood in humans and has also been characterized in several important viral pathogens by us and others [13-17]. In eukaryotic parasites, this process is not that well described. In *T. gondii*, the RNA triphosphatase responsible for the first step of RNA capping was shown to be essential for transcript homeostasis and survival [18], this process, including RNA N7 methylation, was also described for *T. brucei* [19, 20]. In *T. vaginalis* the bifunctional capping enzyme and cap structures have been characterized [12], however its RNA MTases have remained biochemically and structurally uncharacterized. In this study, we structurally and functionally characterized the RNMT enzymes from *T. vaginalis* (hereafter referred to as *Tv*RNMT) and we describe the discovery of submicromolar inhibitors of *Tv*RNMT.

## Results

### Crystal structure of TvRNMT

We aimed to functionally and structurally characterize the RNA cap methylation process in *T. vaginalis*. This parasite encodes two homologues of the N7 RNA MTase, *Tv*RNMT1 (UniProt ID: A2FYD5) and *Tv*RNMT2 (UniProt ID: A2E7Y7). Therefore, we cloned both genes and produced the corresponding recombinant proteins in *E. coli*. We obtained both proteins in soluble form and at high purity (**SI Figure 1**).

We then proceeded with crystallization screening. The proteins were supplemented with their natural ligand SAH at a 2 mM concentration and initial screening for crystals was performed. Initial crystallization trials yielded *Tv*RNMT1 crystals that were of poor quality. Optimization by iterative microseeding produced well-diffracting crystals, from which a complete dataset was collected to 1.6 Å resolution. The crystals belonged to space group P2_1_ and contained one TvRNMT1 molecule in the asymmetric unit. Molecular replacement followed by refinement yielded a high-quality model with Rwork and Rfree values of 19.2% and 22.3%, respectively (SI Table 1).

The structure revealed a Rossmann-like fold typical of MTases, characterized by a central β-sheet (composed here of strands β4, β5, β9 and β10) surrounded by α-helices (α1, α2, α3, α5, α6, α9 and α12), which together form the SAM/SAH-binding site (**Figure 1**). Electron density corresponding to SAH was immediately visible upon molecular replacement within the SAM/SAH-binding pocket formed by these structural elements (**Figure 1B**).

**Fig. 1.**
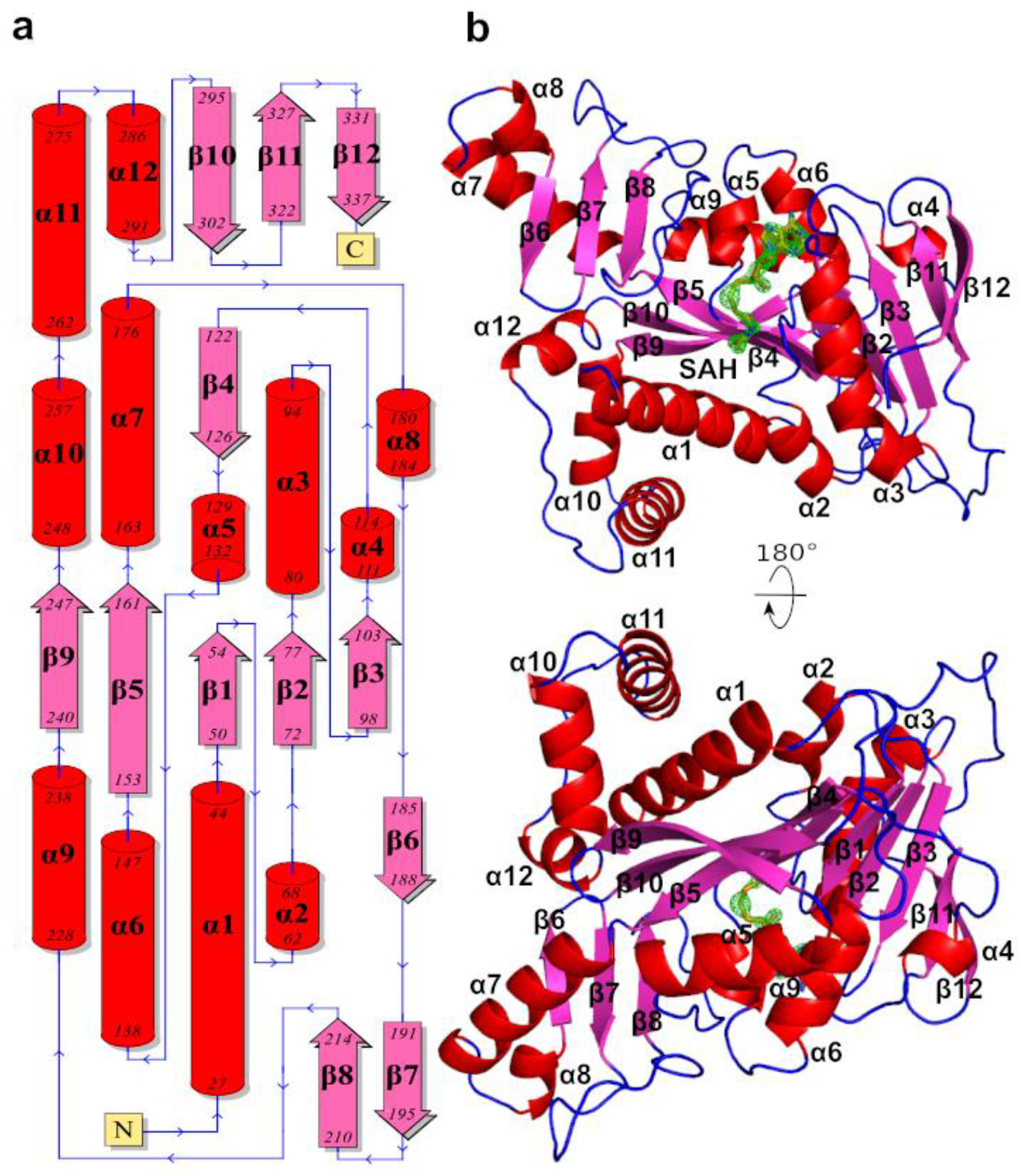
Overall structure and topology of the *T. vaginalis* RNMT. **a**) Topology plot of *Tv*RNMT1 showing the arrangement of α-helices (red cylinders) and β-strands (magenta arrows). Secondary-structure elements are labelled sequentially, and residue numbers corresponding to the boundaries of individual elements are indicated. **b**) Cartoon representation of the *Tv*RNMT structure in two orientations related by a 180° rotation. SAH is shown in stick representation. The Fo-Fc omit map is shown in a green mesh and contoured at 2.5σ.

### Biochemical analysis and screening for inhibitors

In parallel with the crystallographic experiments, we performed a biochemical characterization of both homologues of this enzyme. Cap-RNA MTases differ considerably in their substrate requirements. In principle, relatively long RNAs (tens of nucleotides in length) containing a pre-cap structure represent universal substrates; however, their preparation is tedious and these RNAs are prone to degradation. For some cap-RNA MTases, such substrates are required, as is the case for flaviviral MTases [21, 22], whereas other enzymes accept simpler pre-cap analogues such as GpppG or GpppG-N_4-10_ [23-25]. We therefore tested the substrate requirements of *Tv*RNMT1 and *Tv*RNMT2 and observed that both homologues were active *in vitro* and preferred pre-capped RNA (GpppG/A-N_35_ was used) which was in subsequent experiments used as a substrate. *Tv*RNMT1 was somewhat more active *in vitro* and also accepted GpppG as a substrate (**Figure 2**).

**Figure 2:**
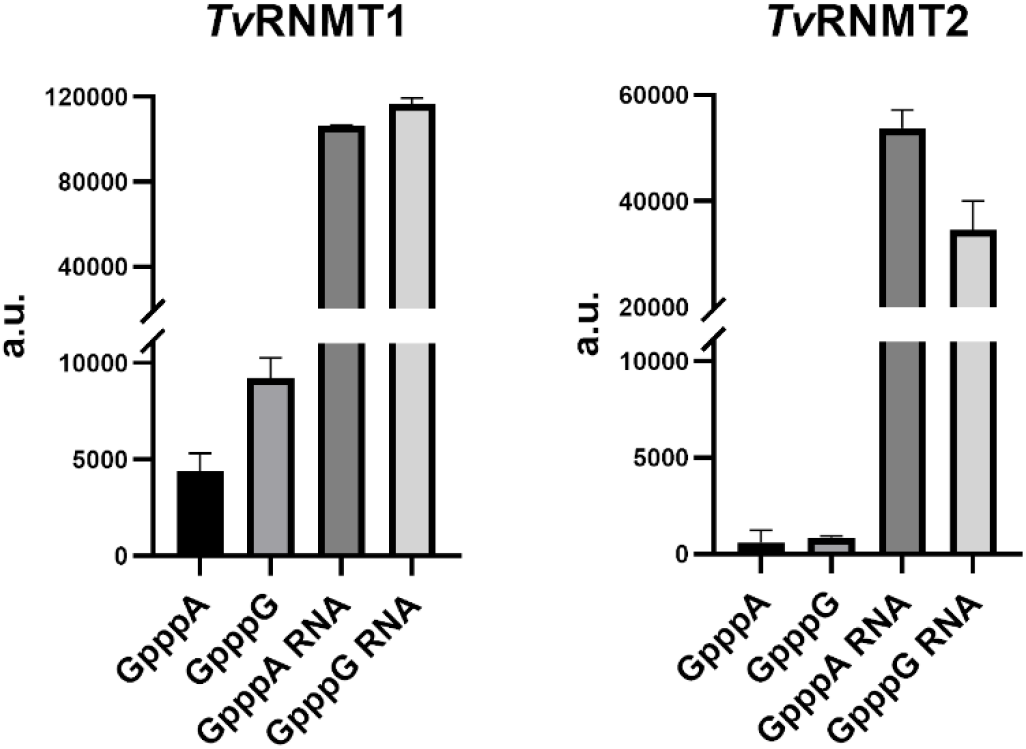
The MTase activity of *Tv*RNMT1 and *Tv*RNMT2 measured with various substrates: GpppA, GpppG, GpppA-capped RNA and GpppG-capped RNA. Measurements were performed in triplicate. *Tv*RNMT2 methylated only capped RNAs, whereas *Tv*RNMT1 also accepted GpppA and GpppG cap analogues.

Next, we screened our small in-house library of SAH analogues to identify *Tv*RNMT inhibitors; the pan-MTase inhibitor sinefungin was included as a positive control. Screening was performed using a universal mass spectrometry-based assay developed previously [26]. This method exploits the fact that, during the methylation reaction, one SAM molecule serves as the methyl donor, resulting in substrate methylation and the formation of one SAH molecule per transferred methyl group. The amount of SAH produced is then quantified.

Several compounds inhibiting *Tv*RNMT were identified, and the most potent compounds are shown (**Figure 3**). Chemically, all of them are SAH analogues, bearing a modified adenine ring. The TO compounds shown in Figure 3 are 7-deaza-SAH analogues bearing bulky aromatic substituents at the C7 position of the pseudoadenine ring. The STM compounds bear a benzylcarboxylate or a (di)benzylcarboxamide group at the same position. Interestingly, the OBO compound is a modified SAH (not 7-deaza), with a dimethylvinyl group at the C-8 position.

**Figure 3:**
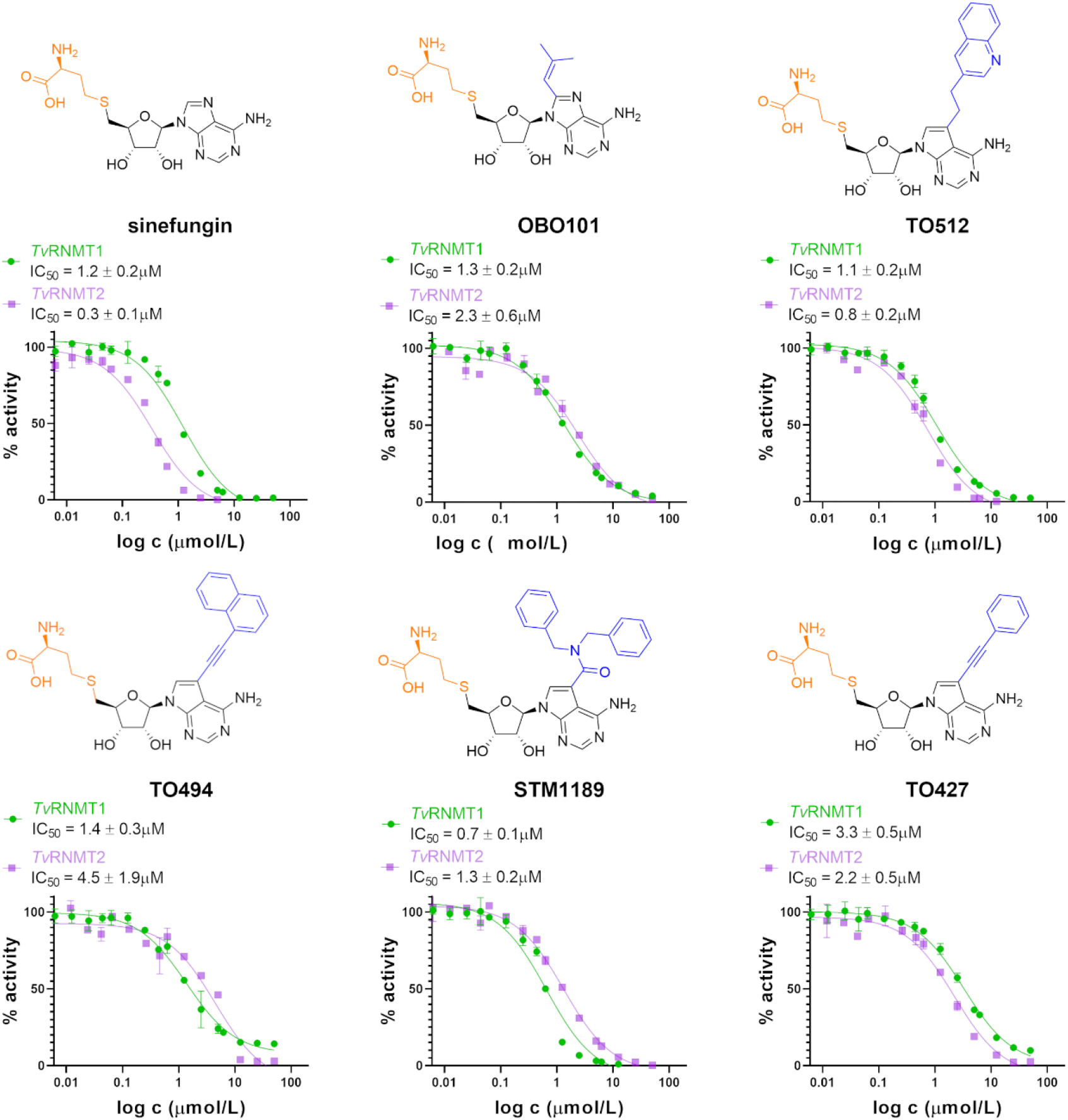
Small-molecule inhibitors of *Tv*RNMT1 and *Tv*RNMT2 and corresponding IC_50_ values. The chemical structures of sinefungin and a series of small-molecule inhibitors examined in this study are shown, together with their respective IC_50_ values and inhibition curves obtained from methyltransferase inhibition assays. Data points and IC_50_ values are reported as the mean ± SEM from triplicate measurements

### The architecture of the SAM/SAH-binding site and inhibitor recognition

To gain structural insight into ligand recognition, we sought to determine crystal structures of *Tv*RNMT in complex with inhibitors. We succeeded in obtaining structures of *Tv*RNMT bound to the pan-methyltransferase inhibitor sinefungin as well as to the compound OBO101 (**SI Figure 2**). These structures enabled a detailed comparison of the binding modes of SAH, sinefungin and OBO101 at atomic resolution (**Figure 4**).

**Figure 4:**
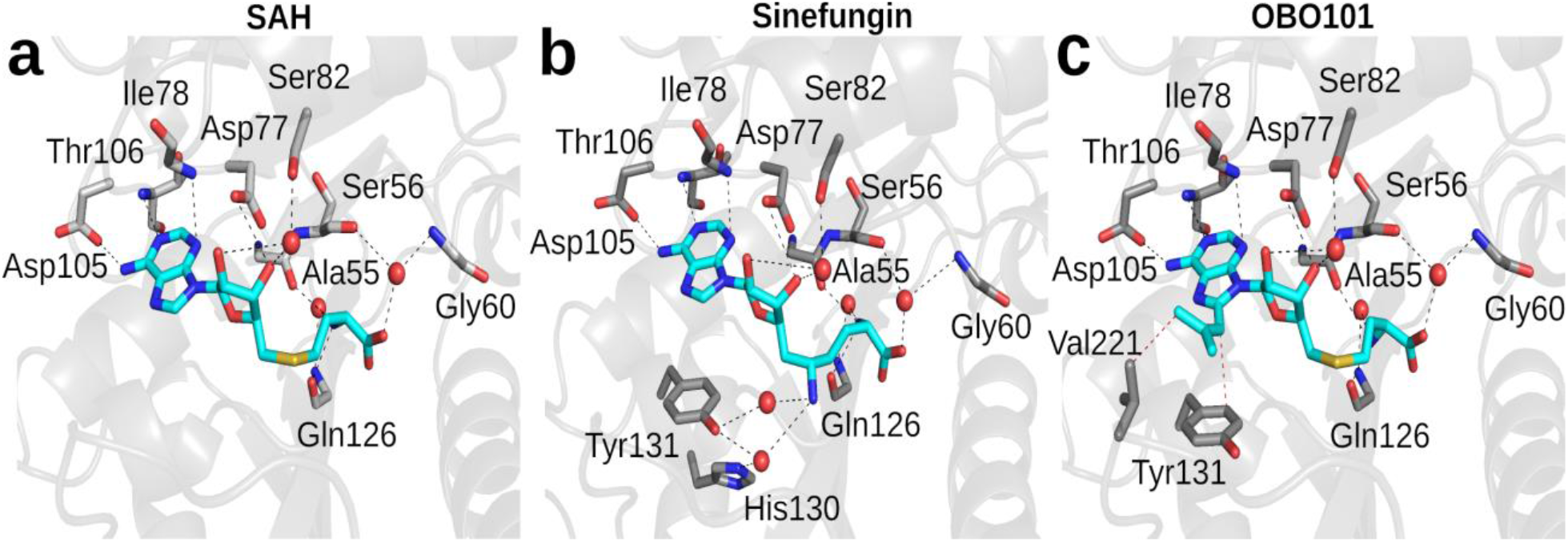
Structural basis of ligand recognition by *Tv*RNMT. (**a–c**) Close-up views of the *Tv*RNMT active site showing the interactions with SAH (**a**), sinefungin (**b**) and OBO101 (**c**). Ligands are shown as sticks with carbon atoms coloured cyan, whereas carbon atoms in protein residues are coloured grey. Water molecules are depicted as red spheres. Hydrogen bonds are indicated by black dashed lines, whereas hydrophobic contacts are shown as red dashed lines. The protein backbone is displayed as a semi-transparent cartoon in light grey. Residues participating in ligand binding are labelled.

All three compounds are bound by a complex network of hydrogen bonds and water bridges. The adenine moiety is accommodated within a predominantly polar pocket and is recognized by residues Asp77, Ser82, Asp105 and Thr106. OBO101 possesses a modified adenine ring bearing a dimethylvinyl moiety at position 8. This moiety effectively exploits the Val221 region; it forms hydrophobic contacts with Val221 and Tyr131 (**Figure 4**), explaining the inhibitory activity of OBO101.

The ribose ring of each compound forms a hydrogen bond with Asp77 and is further stabilised by a water bridge to Ser82. The amino acid portion of these compounds extends toward the solvent-exposed region of the active site and forms direct hydrogen bonds with Gln126 and Ala55, while Ser56 and Gly60 bind the carboxyl group via water bridges. Compared with SAH and OBO101, sinefungin has an additional amino group within its ornithine-like side chain that forms extra polar interactions. Specifically, this amino group forms water bridges with His130 and Tyr131, explaining the inhibitory activity of sinefungin.

### RNA recognition

We performed molecular dynamics (MD) simulations to model the binding of capped RNA into the *Tv*RNMT1 active site. The simulations revealed that the RNA binds to a deep cleft, characterized by highly positively charged residues. The cleft itself is defined by helices α1, α2, α3, α5, α11 and α12 together with strands β1, β2, β4, β5, β8, and β10, as shown in Figure 5. The guanine recognition pocket includes aromatic residues Tyr131 and Phe127 that form π-stacking interactions with the guanine ring. Additionally, the model suggests the 5’-5’ triphosphate bridge is stabilised by the positively charged residues Lys37 and Arg30, which appear to be particularly important for RNA recognition. Overall, the model suggests that *Tv*RNMT recognizes capped RNA through a combination of cap recognition near the active site and non-sequence-specific interactions with the RNA backbone (**Figure 5, SI Figure 3**).

**Figure 5:**
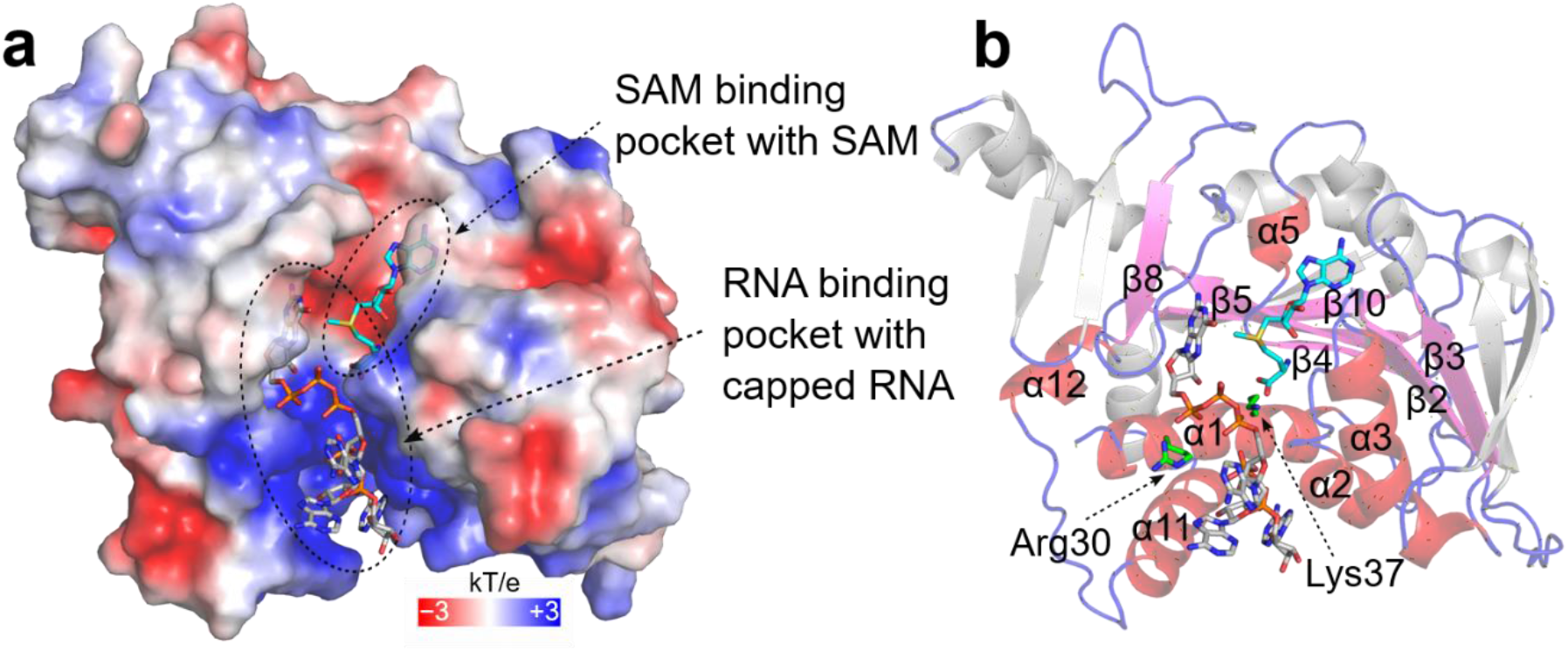
Recognition of capped RNA by *Tv*RNMT. (**a**) A representative model obtained from MD simulations. The TvRNMT1 protein is shown as semitransparent surface and coloured according to the electrostatic surface potential, with negative potential shown in red and positive potential in blue. The SAM molecule as well as the GpppG-capped RNA are shown in stick representation. (**b**) Model of RNA recognition by TvRNMT shown in a cartoon representation. The RNA-binding cleft is formed by highlighted helices and β-strands. Key residues involved in RNA recognition are shown as green sticks.

## Materials and Methods

### Protein expression and purification

The genes encoding *Tv*RNMT1 (UniProt ID: A2FYD5) and *Tv*RNMT2 (UniProt ID: A2E7Y7) were codon-optimized for expression in *E. coli* and commercially synthesized (Azenta/Genewiz). Genes were subcloned into a homemade pSUMO vector (His_8x_-SUMO tag) [27]. Sequences were verified and expression plasmids were transformed into *E. coli* T7 Express lysY/I^q^ cells. Target proteins were expressed in LB medium supplemented with 100 μg/ml ampicillin and grown at 37 °C until the cells reached an optical density of 0.8. Protein expression was initiated by the addition of IPTG (isopropyl-β-D-thiogalactopyranoside) to a final concentration of 250 μM and the cultures were incubated overnight at 18 °C.

The recombinant proteins were purified using our standard protocols for recombinant MTases [17, 21]. Briefly, the bacterial cells were pelleted by centrifugation, resuspended, and sonicated in lysis buffer (20 mM Tris-HCl pH 8, 300 mM NaCl, 20 mM imidazole, 5% glycerol and 2 mM β-mercaptoethanol). The clarified supernatant was loaded onto a Ni-NTA affinity column (Thermo Fisher Scientific) and incubated with the agarose resin for 1 h at 4 °C. After five wash steps with wash buffer (20 mM Tris-HCl pH 8, 1.5 M NaCl, 40 mM imidazole, and 2 mM β-mercaptoethanol) the recombinant proteins were eluted with 25 mL of elution buffer (20 mM Tris-HCl pH 8, 200 mM NaCl, 400 mM imidazole, and 2 mM β-mercaptoethanol). The eluate was dialysed against elution buffer (without imidazole) and N-terminal solubility His_8x_-SUMO tag was cleaved by Ulp1 protease at 4 °C overnight. The cleaved tag was removed by a second round of affinity chromatography and the protein was concentrated and loaded onto a HiLoad 16/600 Superdex 75 gel filtration column (Cytiva), equilibrated with the SEC buffer (10 mM Tris-HCl pH 8, 200 mM NaCl, and 1 mM TCEP (tris(2-carboxyethyl)phosphine)). The peak fractions were collected and concentrated to 20 mg/mL for *Tv*RNMT1 and to 14 mg/mL for *Tv*RNMT2 and directly used for crystallization screening.

### Crystallization, structure determination and refinement

*Tv*RNMT1 and *Tv*RNMT2 proteins were diluted in SEC buffer to a final concentration of 10 mg/mL and supplemented with 2 mM SAH. The crystallization trials were set up using a Mosquito robot (SPT Labtech) in 400 nL drops (mixing protein and reservoir solutions at a 1:1 ratio). Only poor-quality crystals were obtained in the initial experiments. Diffraction-quality crystals were generated by several rounds of seeding under conditions containing 0.2 M magnesium nitrate and 20% (w/v) PEG 3350 for *Tv*RNMT1, and the crystals appeared within 5 days at 18 °C. In the case of *Tv*RNMT2, no crystals were obtained during crystallization trials. Rod-shaped crystals of *Tv*RNMT1 were cryoprotected in well solution supplemented with 20% (v/v) glycerol and plunged into liquid nitrogen.

The crystals with the pan-MTase inhibitor sinefungin and our selected inhibitors were generated by soaking *Tv*RNMT1-SAH crystals with 2 mM sinefungin or 4 mM inhibitor in fresh crystallization solution with the PEG 3350 concentration increased to 25% (w/v).

Crystallographic data collection was performed at the BESSY II electron storage ring (Helmholtz-Zentrum Berlin) [28]. Single rod-shaped crystals were subjected to an X-ray beam with a wavelength of 0.9184 Å at a temperature of 100 K. Data were indexed, integrated, and scaled using the program XDS [29]. The phase problem was solved by molecular replacement in Phaser-MR (Phenix GUI) [30] using a human RNA MTase (PDB ID: 8Q69) [31] as a search model. The initial models were further manually built in COOT [32] and iteratively refined using Phenix.refine [33]. The ligand-restraint CIF dictionary was generated using the online Grade Web Server (Global Phasing Ltd.). All structural figures were prepared using the PyMOL Molecular Graphics System v3.0 (Schrödinger, LLC). The Protein-Ligand Interaction Profiler (PLIP) online server tool was used to identify residues involved in protein:ligand interactions in an unbiased manner [34]. Crystallographic data statistics are summarized in SI Table 1.

### Preparation of capped RNA for MTase assay of TvRNMT2

To generate substrates for the MTase assays, 35-nucleotide GpppA-capped and GpppG-capped RNAs were prepared by in vitro transcription using T7 RNA polymerase. The transcription reaction employed two annealed oligonucleotides: 5’-CAGTAATACGACTCACTATAGGGGAAGCGGGCATGCGGCCAGCCATAGCCGATC-3’ 5’-TGATCGGCTATGGCTGGCCGCATGCCCGCTTCCCCTATAGTGAGTCGTATTACTG-3’

These oligonucleotides were hybridized by gradually cooling the mixture from 95 °C to 20 °C over 40 min in a buffer containing 10 mM Tris-HCl (pH 8.0), 50 mM NaCl, and 1 mM EDTA. This produced a double-stranded DNA template comprising a T7 promoter sequence (CAGTAATACGACTCACTATAG) upstream of the RNA coding region (GGGGAAGCGGGCATGCGGCCAGCCATAGCCGATCA). Transcription was carried out using the TranscriptAid T7 High Yield Transcription Kit (Thermo Fisher Scientific) in the presence of either the GpppA or the GpppG cap analogue. The resulting capped RNAs (GpppA-RNA and GpppG-RNA) were purified by phenol–chloroform extraction and stored at −20 °C until use.

### Methyltransferase activity assay

The MTase activities of *Tv*RNMT1 and *Tv*RNMT2 were initially tested using various substrates: GpppA, GpppG, GpppA-capped RNA, and GpppG-capped RNA. The 4 µL reaction mixture contained 0.5 µM *Tv*RNMT1 or *Tv*RNMT2, reaction buffer (5 mM Tris-HCl, pH 8.0; 1 mM TCEP; 0.1 mg/mL BSA; 1 mM MgCl_2_), 8 µM S-adenosyl-L-methionine (SAM; BLDpharm), and 8 µM substrate. Reactions were incubated for 120 min at 25 °C and quenched by the addition of formic acid to a final concentration of 5%.

Samples were analysed using an Echo system coupled to a Sciex 6500 triple-quadrupole mass spectrometer equipped with an electrospray ionization source. MTase activity was quantified based on the formation of S-adenosylhomocysteine (SAH). The instrument was operated in multiple reaction monitoring (MRM) mode with an interface temperature of 350 °C. Instrument settings were as follows: declustering potential, 20 V; entrance potential, 10 V; and collision energy, 28 eV. A 10 nL injection volume was analysed in 100% methanol at a flow rate of 0.40 mL/min. The transition m/z 385.1 → 134.1 corresponding to SAH was used for quantification.

For *Tv*RNMT1 inhibition assays, compounds were added either at 10 µM for screening or in the range of 0 - 50 µM for IC_50_ determination. *Tv*RNMT1 activity was determined by measuring the conversion of GpppA-capped RNA to m^7^GpppA-capped RNA. The 4 µL reaction mixture contained 0.4 µM *Tv*RNMT1, reaction buffer (5 mM Tris-HCl, pH 8.0; 1 mM TCEP; 0.1 mg/mL BSA, 1 mM MgCl_2_), 8 µM SAM, and 4 µM GpppA-capped RNA. DB RNase inhibitor (DIANA Biotechnologies) was included in all reactions. Reactions were incubated for 60 min at 25 °C, quenched with formic acid to a final concentration of 5%, and analysed using the Echo system as described above.

Inhibition assays for *Tv*RNMT2 were performed the same way as for *Tv*RNMT1, but the reaction mixture contained 0.5 µM *Tv*RNMT2 (instead of 0.4 µM *Tv*RNMT1) and 6 µM GpppA-capped RNA (instead of 4 µM GpppA-capped RNA).

### Organic synthesis

The TO and STM compounds had been synthesized previously [26, 35, 36] and were available in our laboratory. OBO101 was synthesized using approaches that are standard in medicinal chemistry, as detailed in the SI Materials and Methods.

### Docking experiments

Molecular docking was performed with GOLD using the ChemPLP scoring function. Ligand structures were prepared in ACD/ChemSketch 2023.1.2, converted to MOL2 format using Open Babel, and geometry-optimized by the PM7 semiempirical method in MOPAC 23.2.5, following the protocol used in our previous studies. The protein coordinates were taken from our newly determined crystal structure and prepared for docking in PDB/MOL2 format, while the ligands were treated as flexible. The binding site was defined from protein atoms surrounding the reference ligand, using a cavity radius of 6 Å. A structurally related reference ligand was used to apply a similarity constraint and preserve the experimentally observed binding orientation. Six crystallographic water molecules, H_2_O-1, H_2_O-2, H_2_O-10, H_2_O-57, H_2_O-73, and H_2_O-147, were retained and treated as toggle/spin waters to account for possible water-mediated interactions. For each ligand, 10 independent docking runs were performed using automatic GOLD genetic algorithm settings; poses were clustered with an RMSD tolerance of 1.5 Å, and the top three solutions were retained, ranked by ChemPLP score, and visually inspected for consistency with the reference binding mode and key protein–ligand interactions.

### Molecular dynamics (MD) simulations

The simulation system comprised RNMT1 in complex with SAM and a non-methylated, capped three-nucleotide RNA. The capped RNA coordinates were derived from the structure of the mpox MTase VP39 [26], and the starting model of the RNMT1/RNA complex was generated using AlphaFold/Boltz 3 [37, 38]. The MD simulations were set up using the input generator on the CHARMM-GUI web server [39, 40].

The *Tv*RNMT1/RNA complex was centred in a cubic simulation cell with an edge length of 8.7 nm and immersed in explicit TIP3P water. Na^+^ and Cl^−^ ions were introduced both to neutralize the total charge and to obtain a final salt concentration of 150 mM. Protein and RNA were described using the CHARMM36m and CHARMM36 force fields, respectively [41-43]. Parameters assigned to guanosine triphosphate were used to describe the triphosphate bridge of the RNA cap, whereas SAH and SAM were parameterized with the CHARMM General Force Field [44].

The MD simulations were carried out using the Nanoscale Molecular Dynamics (NAMD) software package [45] under periodic boundary conditions and a 2 fs integration time step. Short-range non-bonded interactions were smoothly switched off between 1.0 and 1.2 nm. Long-range electrostatics were evaluated with the particle-mesh Ewald method using a grid spacing of 0.1 nm. Each of the two systems was energy-minimized using the conjugate-gradient method and then equilibrated using a standard procedure using input files obtained from the CHARMM-GUI input generator. The temperature was maintained at 30 °C with a Langevin thermostat using a damping coefficient of 1 ps^-1^. During production simulations, the pressure was held at 1 atm using the Langevin piston Nosé–Hoover method, with a damping timescale of 25 fs and an oscillation period of 50 fs. Soft harmonic restraints with a force constant of 100 kcal mol^-1^ nm^-2^ were imposed on the protein backbone atoms and on the non-hydrogen atoms of SAH/SAM. Protein side-chain atoms and RNA atoms were unrestrained so that the RNA could adapt to the RNMT1 crystal structure. Two independent 100-ns production runs were performed, and the MD trajectories were inspected and analysed using VMD [46].

## Discussion

In this study, we structurally and functionally characterized N7 RNA cap methylation in *T. vaginalis*. Biochemical characterization demonstrated that both *Tv*RNMT homologues are active in vitro and prefer capped RNA substrates. Furthermore, screening of our in-house library of SAH analogues identified several compounds that inhibited *Tv*RNMTs with low-micromolar to submicromolar IC_50_ values. STM1189 was more potent than the pan-MTase inhibitor sinefungin against *Tv*RNMT1, whereas sinefungin remained the most potent inhibitor of TvRNMT2. We solved the crystal structure of *Tv*RNMT1 in complex with SAH and determined additional structures with sinefungin and the inhibitor OBO101, providing detailed insights into ligand recognition and inhibitor binding. Finally, structural modelling suggested a plausible mode of capped-RNA recognition and identified positively charged residues involved in RNA recognition.

The crystal structures revealed the binding modes of the pan-MTase inhibitor sinefungin and the compound OBO101. Unfortunately, we were unable to obtain crystal structures of *Tv*RNMT1 bound to the TO compounds despite extensive efforts. One possible explanation is that the bulky C7 substituents of the TO compounds are incompatible with the packing of the *Tv*RNMT1 crystal form. Molecular docking was therefore used to evaluate the binding mode of the SAH-derived ligands in the active site of the newly obtained crystal structure. The adenine/ribose region remained positioned in the conserved part of the binding site and the amino acid-derived portion of the ligands was oriented toward the polar region involved in recognition of the SAH scaffold. Inclusion of selected crystallographic water molecules allowed retention of potential water-mediated contacts, which may be important for stabilizing the ligand-binding mode. The docked compounds generally adopted poses consistent with the reference ligand, indicating that the applied similarity constraint preserved the characteristic SAH-like orientation in the binding pocket. This analysis revealed that the aromatic moieties of the TO compounds efficiently exploit the Val221 region, similarly to the dimethylvinyl group of OBO101 (**Figure 6**). In addition to Val221, residues Lys138, Tyr224 and Tyr131 also play important roles in inhibitor binding.

**Figure 6:**
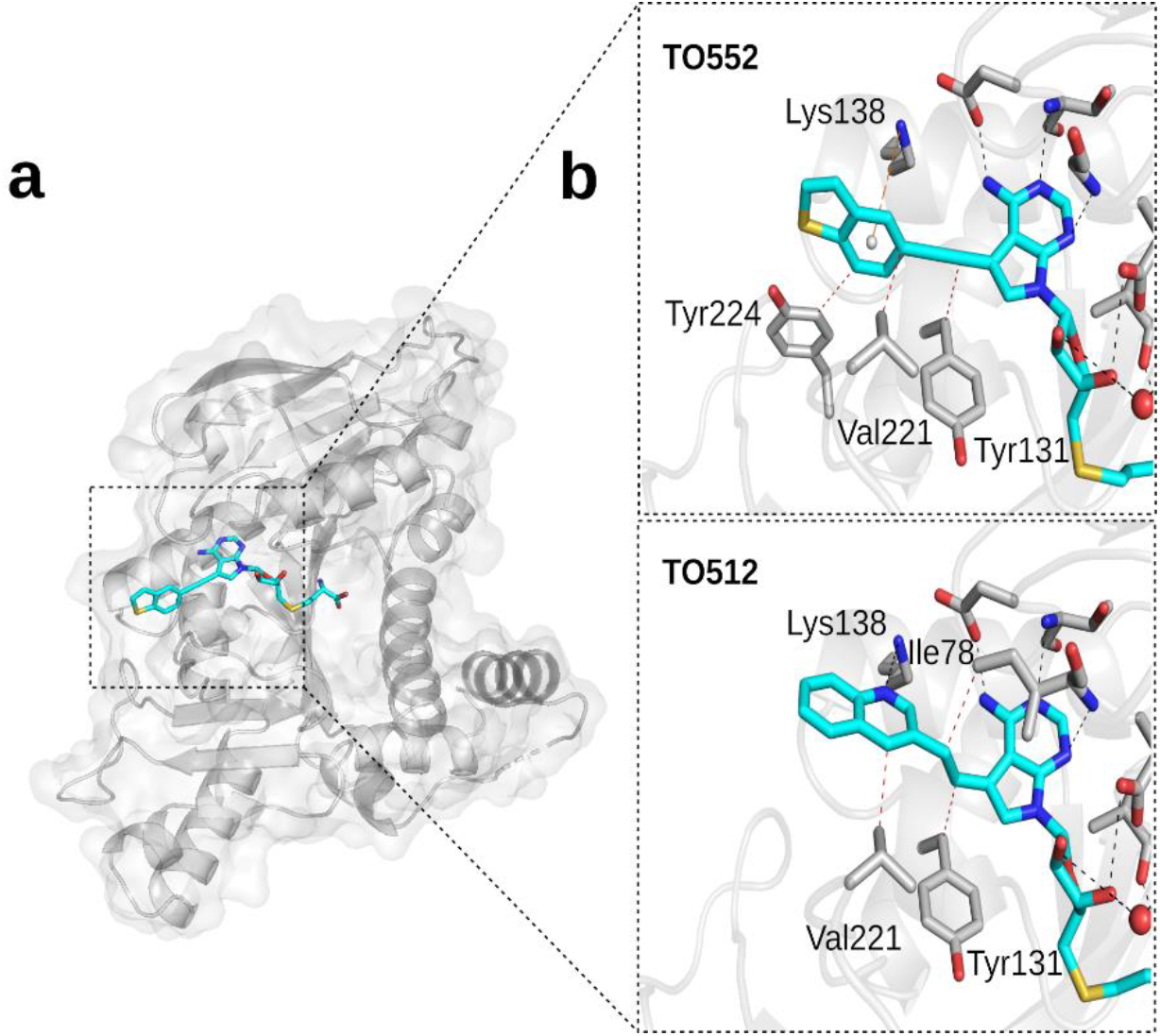
Binding mode of the TO compounds to *Tv*RNMT. Key amino acid residues of *Tv*RNMT1 involved in inhibitor binding are shown in stick representation. Ligands are depicted in cyan with standard heteroatom colouring. Hydrogen bonds are shown as dashed black lines, hydrophobic interactions as red dashed lines, and π–cation interactions as orange dashed lines.

The binding mode of the inhibitors is partially conserved across RNA cap MTases from pathogenic organisms. Several of the compounds (TO427 and TO494) also inhibit the RNA cap MTase from the important pathogens mpox virus and SARS-CoV-2 [26, 35]. As shown previously in cells, compounds bearing large aromatic substituents at the 7-deaza position showed little or no detectable cytotoxicity, whereas several analogues bearing smaller substituents were cytotoxic [47]. From a structural perspective, it seems that the human RNA capping enzyme, *Hs*RNMT, cannot accommodate compounds bearing a pseudoadenine ring bearing such a large substituent. Altogether, our results suggest that targeting the RNA cap methylation pathway is feasible in *T. vaginalis*. Although *T. vaginalis* possesses two homologues of the N7 RNA cap MTase, inhibitors can be developed to target both homologues, an effect that was especially pronounced for the compounds shown in Figure 3.

## Limitations of the study

Although this study identifies inhibitors of *Tv*RNMTs and defines their binding modes at atomic resolution, their therapeutic potential remains to be established. Assessment of membrane permeability, metabolic stability, pharmacokinetics, tissue distribution and toxicity will be required to determine their suitability for drug development.

## Use of artificial intelligence

ChatGPT (GPT-5.6 Sol, OpenAI) was used solely for language editing to improve the clarity, grammar and style of the manuscript. It was not used for data analysis, data interpretation, generation of scientific conclusions or preparation of figures. All scientific content and conclusions were generated and verified by the authors.

## Acknowledgments

We are grateful to Petr Pachl for help with X-ray data collection. We thank the Helmholtz-Zentrum Berlin für Materialien und Energie and the EMBL Hamburg at the PETRA III storage ring (DESY, Hamburg, Germany) for the allocation of synchrotron radiation beamtime. The molecular dynamics simulations were carried out using the supercomputer resources at the Centre of Informatics - Tricity Academic Supercomputer Network (CI TASK) in Gdansk, Poland.

## Funding

This work was supported by the Czech Science Foundation (GACR), grant No. 26-20827S “RNA Capping Enzymes in T. vaginalis: Structural and Functional Characterization for Drug Discovery”.

## Competing interests

The authors declare no competing interests.

## Contributions

J.B., D.C., M.S., O.B., T.O., and M.D. performed experiments. B.R. performed computer simulations. R.N. performed docking experiments. M.D., R.N. and E.B. supervised the project. E.B. conceived the project and obtained funding. J.B., M.D., B.R., R.N. and E.B. wrote the manuscript.

## Notes

### Competing Interest Statement

The authors have declared no competing interest.

## References

1. Munoz, C., J. San Francisco, B. Gutierrez, and J. Gonzalez, Role of the Ubiquitin-Proteasome Systems in the Biology and Virulence of Protozoan Parasites. Biomed Res Int, 2015. 2015: p. 141526.

2. Edwards, T., P. Burke, H. Smalley, and G. Hobbs, Trichomonas vaginalis: Clinical relevance, pathogenicity and diagnosis. Critical Reviews in Microbiology, 2016. 42(3): p. 406–417.

3. Rowley, J., S. Vander Hoorn, E. Korenromp, N. Low, M. Unemo, L.J. Abu-Raddad, R.M. Chico, A. Smolak, L. Newman, S. Gottlieb, S.S. Thwin, N. Broutet, and M.M. Taylor, Chlamydia, gonorrhoea, trichomoniasis and syphilis: global prevalence and incidence estimates, 2016. Bulletin of the World Health Organization, 2019. 97(8): p. 548–+.

4. Alsaad, R.K.A., Past, present and future of Trichomonas vaginalis: a review study. Ann Parasitol, 2022. 68(3): p. 409–419.

5. Bouchemal, K., C. Bories, and P.M. Loiseau, Strategies for Prevention and Treatment of Trichomonas vaginalis Infections. Clin Microbiol Rev, 2017. 30(3): p. 811–825.

6. Van Gerwen, O.T., A.F. Camino, J. Sharma, P.J. Kissinger, and C.A. Muzny, Epidemiology, Natural History, Diagnosis, and Treatment of Trichomonas vaginalis in Men. Clin Infect Dis, 2021. 73(6): p. 1119–1124.

7. Alessio, C. and P. Nyirjesy, Management of Resistant Trichomoniasis. Curr Infect Dis Rep, 2019. 21(9): p. 31.

8. Marques-Silva, M., C. Lisboa, N. Gomes, and A.G. Rodrigues, Trichomonas vaginalis and growing concern over drug resistance: a systematic review. J Eur Acad Dermatol Venereol, 2021. 35(10): p. 2007–2021.

9. Ramanathan, A., G.B. Robb, and S.H. Chan, mRNA capping: biological functions and applications. Nucleic Acids Res, 2016. 44(16): p. 7511–26.

10. Shuman, S., RNA capping: progress and prospects. RNA, 2015. 21(4): p. 735–7.

11. Cowling, V.H., Regulation of mRNA cap methylation. Biochem J, 2009. 425(2): p. 295–302.

12. Simoes-Barbosa, A., R.P. Hirt, and P.J. Johnson, A Metazoan/Plant-like Capping Enzyme and Cap Modified Nucleotides in the Unicellular Eukaryote. Plos Pathogens, 2010. 6(7).

13. Krafcikova, P., J. Silhan, R. Nencka, and E. Boura, Structural analysis of the SARS-CoV-2 methyltransferase complex involved in RNA cap creation bound to sinefungin. Nat Commun, 2020. 11(1): p. 3717.

14. Skvara, P., D. Chalupska, M. Klima, J. Kozic, J. Silhan, and E. Boura, Structural basis for RNA-cap recognition and methylation by the mpox methyltransferase VP39. Antiviral Res, 2023. 216: p. 105663.

15. Decroly, E., C. Debarnot, F. Ferron, M. Bouvet, B. Coutard, I. Imbert, L. Gluais, N. Papageorgiou, A. Sharff, G. Bricogne, M. Ortiz-Lombardia, J. Lescar, and B. Canard, Crystal structure and functional analysis of the SARS-coronavirus RNA cap 2’-O-methyltransferase nsp10/nsp16 complex. PLoS Pathog, 2011. 7(5): p. e1002059.

16. Coutard, B., K. Barral, J. Lichiere, B. Selisko, B. Martin, W. Aouadi, M.O. Lombardia, F. Debart, J.J. Vasseur, J.C. Guillemot, B. Canard, and E. Decroly, Zika Virus Methyltransferase: Structure and Functions for Drug Design Perspectives. Journal of Virology, 2017. 91(5).

17. Klima, M., A. Khalili Yazdi, F. Li, I. Chau, T. Hajian, A. Bolotokova, H.U. Kaniskan, Y. Han, K. Wang, D. Li, M. Luo, J. Jin, E. Boura, and M. Vedadi, Crystal structure of SARS-CoV-2 nsp10-nsp16 in complex with small molecule inhibitors, SS148 and WZ16. Protein Sci, 2022. 31(9): p. e4395.

18. Aswale, K.R., P. Kashyap, and A.S. Deshmukh, RNA triphosphatase-mediated mRNA capping is essential for maintaining transcript homeostasis and the survival of Toxoplasma gondii. Nat Commun, 2025. 16(1): p. 5452.

19. Hall, M.P. and C.K. Ho, Functional characterization of a 48 kDa cap 2 RNA methyltransferase. Nucleic Acids Research, 2006. 34(19): p. 5594–5602.

20. Hall, M.P. and C.K. Ho, Characterization of a RNA cap (guanine N-7) methyltransferase. Rna, 2006. 12(3): p. 488–497.

21. Krejcova, K., P. Krafcikova, M. Klima, D. Chalupska, K. Chalupsky, E. Zilecka, and E. Boura, Structural and functional insights in flavivirus NS5 proteins gained by the structure of Ntaya virus polymerase and methyltransferase. Structure, 2024.

22. Yap, L.J., D. Luo, K.Y. Chung, S.P. Lim, C. Bodenreider, C. Noble, P.Y. Shi, and J. Lescar, Crystal structure of the dengue virus methyltransferase bound to a 5’-capped octameric RNA. PLoS One, 2010. 5(9).

23. Lockless, S.W., H.T. Cheng, A.E. Hodel, F.A. Quiocho, and P.D. Gershon, Recognition of capped RNA substrates by VP39, the vaccinia virus-encoded mRNA cap-specific 2′--methyltransferase. Biochemistry, 1998. 37(23): p. 8564–8574.

24. Benoni, R., P. Krafcikova, M.R. Baranowski, J. Kowalska, E. Boura, and H. Cahova, Substrate Specificity of SARS-CoV-2 Nsp10-Nsp16 Methyltransferase. Viruses, 2021. 13(9): p. 1722.

25. Hausmann, S., S.S. Zheng, C. Fabrega, S.W. Schneller, C.D. Lima, and S. Shuman, mRNA cap (guanine N-7) methyltransferase. Journal of Biological Chemistry, 2005. 280(21): p. 20404–20412.

26. Silhan, J., M. Klima, T. Otava, P. Skvara, D. Chalupska, K. Chalupsky, J. Kozic, R. Nencka, and E. Boura, Discovery and structural characterization of monkeypox virus methyltransferase VP39 inhibitors reveal similarities to SARS-CoV-2 nsp14 methyltransferase. Nat Commun, 2023. 14(1): p. 2259.

27. Dubankova, A. and E. Boura, Structure of the yellow fever NS5 protein reveals conserved drug targets shared among flaviviruses. Antiviral Res, 2019. 169: p. 104536.

28. Mueller, U., T. Barthel, L.S. Benz, V. Bon, T. Crosskey, C.G. Dieguez, R. Förster, C. Gless, T. Hauss, U. Heinemann, M. Hellmig, D. James, F. Lennartz, M. Oelker, R. Ovsyannikov, P. Singh, M.C. Wahl, G. Weber, and M.S. Weiss, The macromolecular crystallography beamlines of the Helmholtz-Zentrum Berlin at the BESSY II storage ring: history, current status and future directions. Journal of Synchrotron Radiation, 2025. 32: p. 766–778.

29. Kabsch, W., Xds. Acta Crystallogr D Biol Crystallogr, 2010. 66(Pt 2): p. 125–32.

30. McCoy, A.J., R.W. Grosse-Kunstleve, P.D. Adams, M.D. Winn, L.C. Storoni, and R.J. Read, Phaser crystallographic software. J Appl Crystallogr, 2007. 40(Pt 4): p. 658–674.

31. Pearson, L.A., A.P. Petit, C. Mendoza Martinez, F. Bellany, D. Lin, S. Niven, R. Swift, T. Eadsforth, P. Fyfe, M. Paul, V. Postis, X. Hu, V.H. Cowling, and D.W. Gray, Characterisation of RNA guanine-7 methyltransferase (RNMT) using a small molecule approach. Biochem J, 2025. 482(4).

32. Emsley, P., B. Lohkamp, W.G. Scott, and K. Cowtan, Features and development of Coot. Acta Crystallogr D Biol Crystallogr, 2010. 66(Pt 4): p. 486–501.

33. Liebschner, D., P.V. Afonine, M.L. Baker, G. Bunkoczi, V.B. Chen, T.I. Croll, B. Hintze, L.W. Hung, S. Jain, A.J. McCoy, N.W. Moriarty, R.D. Oeffner, B.K. Poon, M.G. Prisant, R.J. Read, J.S. Richardson, D.C. Richardson, M.D. Sammito, O.V. Sobolev, D.H. Stockwell, T.C. Terwilliger, A.G. Urzhumtsev, L.L. Videau, C.J. Williams, and P.D. Adams, Macromolecular structure determination using X-rays, neutrons and electrons: recent developments in Phenix. Acta Crystallographica Section D-Structural Biology, 2019. 75: p. 861–877.

34. Schake, P., S.N. Bolz, K. Linnemann, and M. Schroeder, PLIP 2025: introducing protein-protein interactions to the protein-ligand interaction profiler. Nucleic Acids Res, 2025. 53(W1): p. W463–W465.

35. Otava, T., M. Sala, F. Li, J. Fanfrlik, K. Devkota, S. Perveen, I. Chau, P. Pakarian, P. Hobza, M. Vedadi, E. Boura, and R. Nencka, The Structure-Based Design of SARS-CoV-2 nsp14 Methyltransferase Ligands Yields Nanomolar Inhibitors. ACS Infect Dis, 2021. 7(8): p. 2214–2220.

36. Stefek, M., M. Klima, T. Otava, D. Chalupska, M. Dejmek, R. Nencka, and E. Boura, Branched C7-Substituted 7-Deaza-SAH Analogues Occupy the Entire SAM-Binding Pocket of Mpox Virus VP39 and Dengue Virus NS5 Methyltransferases. bioRxiv, 2026: p. 2026.07. 28.741162.

37. Wohlwend, J., G. Corso, S. Passaro, N. Getz, M. Reveiz, K. Leidal, W. Swiderski, L. Atkinson, T. Portnoi, and I. Chinn, Boltz-1 democratizing biomolecular interaction modeling. BioRxiv, 2025: p. 2024.11. 19.624167.

38. Abramson, J., J. Adler, J. Dunger, R. Evans, T. Green, A. Pritzel, O. Ronneberger, L. Willmore, A.J. Ballard, J. Bambrick, S.W. Bodenstein, D.A. Evans, C.C. Hung, M. O’Neill, D. Reiman, K. Tunyasuvunakool, Z. Wu, A. Zemgulyte, E. Arvaniti, C. Beattie, O. Bertolli, A. Bridgland, A. Cherepanov, M. Congreve, A.I. Cowen-Rivers, A. Cowie, M. Figurnov, F.B. Fuchs, H. Gladman, R. Jain, Y.A. Khan, C.M.R. Low, K. Perlin, A. Potapenko, P. Savy, S. Singh, A. Stecula, A. Thillaisundaram, C. Tong, S. Yakneen, E.D. Zhong, M. Zielinski, A. Zidek, V. Bapst, P. Kohli, M. Jaderberg, D. Hassabis, and J.M. Jumper, Accurate structure prediction of biomolecular interactions with AlphaFold 3. Nature, 2024. 630(8016): p. 493–500.

39. Jo, S., T. Kim, V.G. Iyer, and W. Im, CHARMM-GUI: a web-based graphical user interface for CHARMM. J Comput Chem, 2008. 29(11): p. 1859–65.

40. Lee, J., X. Cheng, J.M. Swails, M.S. Yeom, P.K. Eastman, J.A. Lemkul, S. Wei, J. Buckner, J.C. Jeong, Y.F. Qi, S. Jo, V.S. Pande, D.A. Case, C.L. Brooks, A.D. MacKerell, J.B. Klauda, and W. Im, CHARMM-GUI Input Generator for NAMD, GROMACS, AMBER, OpenMM, and CHARMM/OpenMM Simulations Using the CHARMM36 Additive Force Field. Journal of Chemical Theory and Computation, 2016. 12(1): p. 405–413.

41. Huang, J., S. Rauscher, G. Nawrocki, T. Ran, M. Feig, B.L. de Groot, H. Grubmuller, and A.D. MacKerell, Jr., CHARMM36m: an improved force field for folded and intrinsically disordered proteins. Nat Methods, 2017. 14(1): p. 71–73.

42. Huang, J. and A.D. MacKerell, Jr., CHARMM36 all-atom additive protein force field: validation based on comparison to NMR data. J Comput Chem, 2013. 34(25): p. 2135–45.

43. Denning, E.J., U.D. Priyakumar, L. Nilsson, and A.D. Mackerell, Jr., Impact of 2’-hydroxyl sampling on the conformational properties of RNA: update of the CHARMM all-atom additive force field for RNA. J Comput Chem, 2011. 32(9): p. 1929–43.

44. Vanommeslaeghe, K., E. Hatcher, C. Acharya, S. Kundu, S. Zhong, J. Shim, E. Darian, O. Guvench,P. Lopes, I. Vorobyov, and A.D. Mackerell, Jr., CHARMM general force field: A force field for drug-like molecules compatible with the CHARMM all-atom additive biological force fields. J Comput Chem, 2010. 31(4): p. 671–90.

45. Phillips, J.C., D.J. Hardy, J.D.C. Maia, J.E. Stone, J.V. Ribeiro, R.C. Bernardi, R. Buch, G. Fiorin, J. Henin, W. Jiang, R. McGreevy, M.C.R. Melo, B.K. Radak, R.D. Skeel, A. Singharoy, Y. Wang, B. Roux, A. Aksimentiev, Z. Luthey-Schulten, L.V. Kale, K. Schulten, C. Chipot, and E. Tajkhorshid, Scalable molecular dynamics on CPU and GPU architectures with NAMD. Journal of Chemical Physics, 2020. 153(4).

46. Humphrey, W., A. Dalke, and K. Schulten, VMD: visual molecular dynamics. J Mol Graph, 1996. 14(1): p. 33–8, 27-8.

47. Zgarbova, M., T. Otava, J. Silhan, R. Nencka, J. Weber, and E. Boura, Inhibitors of mpox VP39 2’-O methyltransferase efficiently inhibit the monkeypox virus. Antiviral Res, 2023. 218: p. 105714.

