## Supplementary Figures, Tables and Methods for "Characterization of the N7 RNA cap methyltransferase from *Trichomonas vaginalis* and inhibitor discovery"

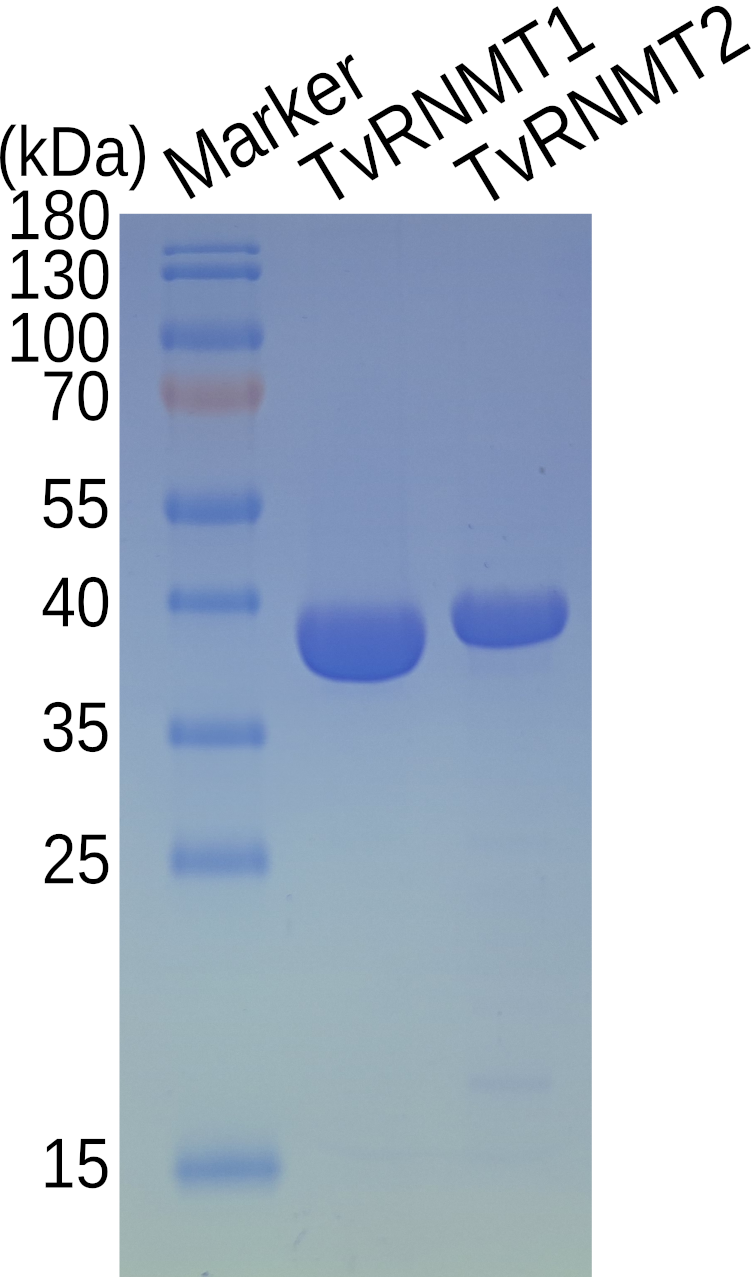

SI Figure 1 – **SDS-PAGE analysis of purified recombinant proteins**. A 15% acrylamide gel was loaded with ~1 µg of each protein. The lanes on the SDS-PAGE gel were as follows: lane 1, protein molecular mass marker, with sizes shown on the left in kilodaltons; lane 2, TvRNMT1, expected molecular mass 40.5 kDa; lane 3, TvRNMT2, expected molecular mass 39.5 kDa.

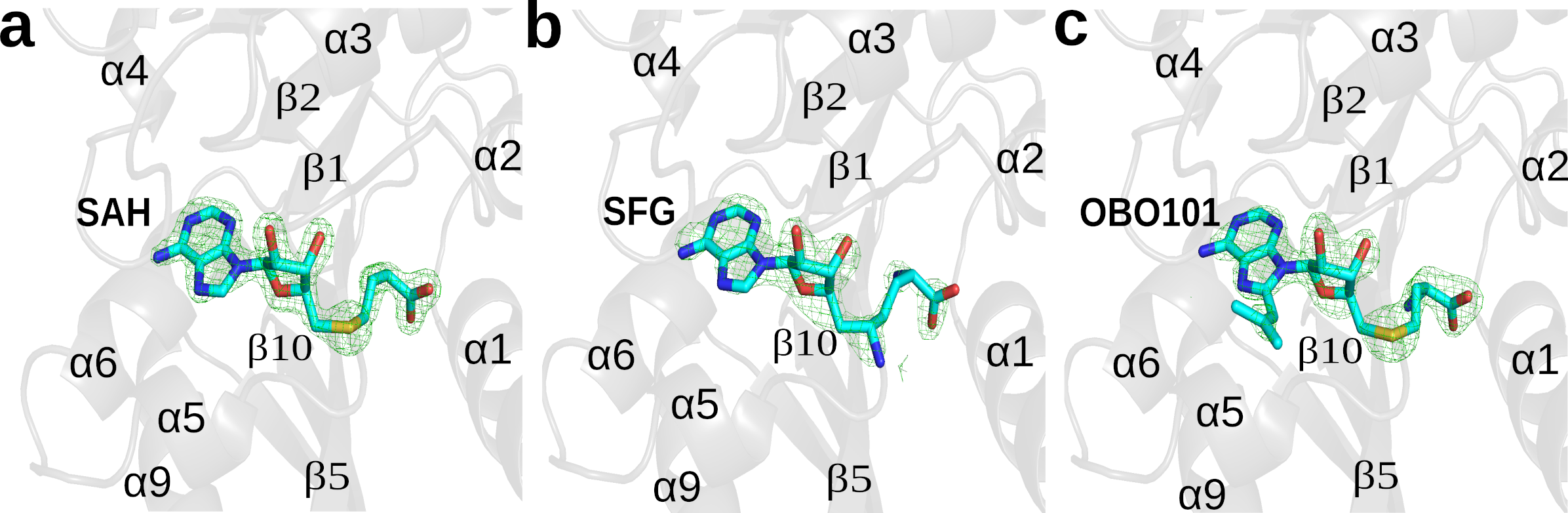

SI Figure 2 – **Electron density maps observed for the ligands.** The Fo–Fc omit map is contoured at 2.5σ and shown as a green mesh. The protein is shown in cartoon representation and coloured grey. Ligands are shown in stick representation with standard colouring for heteroatoms (sulfur, yellow; nitrogen, blue; oxygen, red).

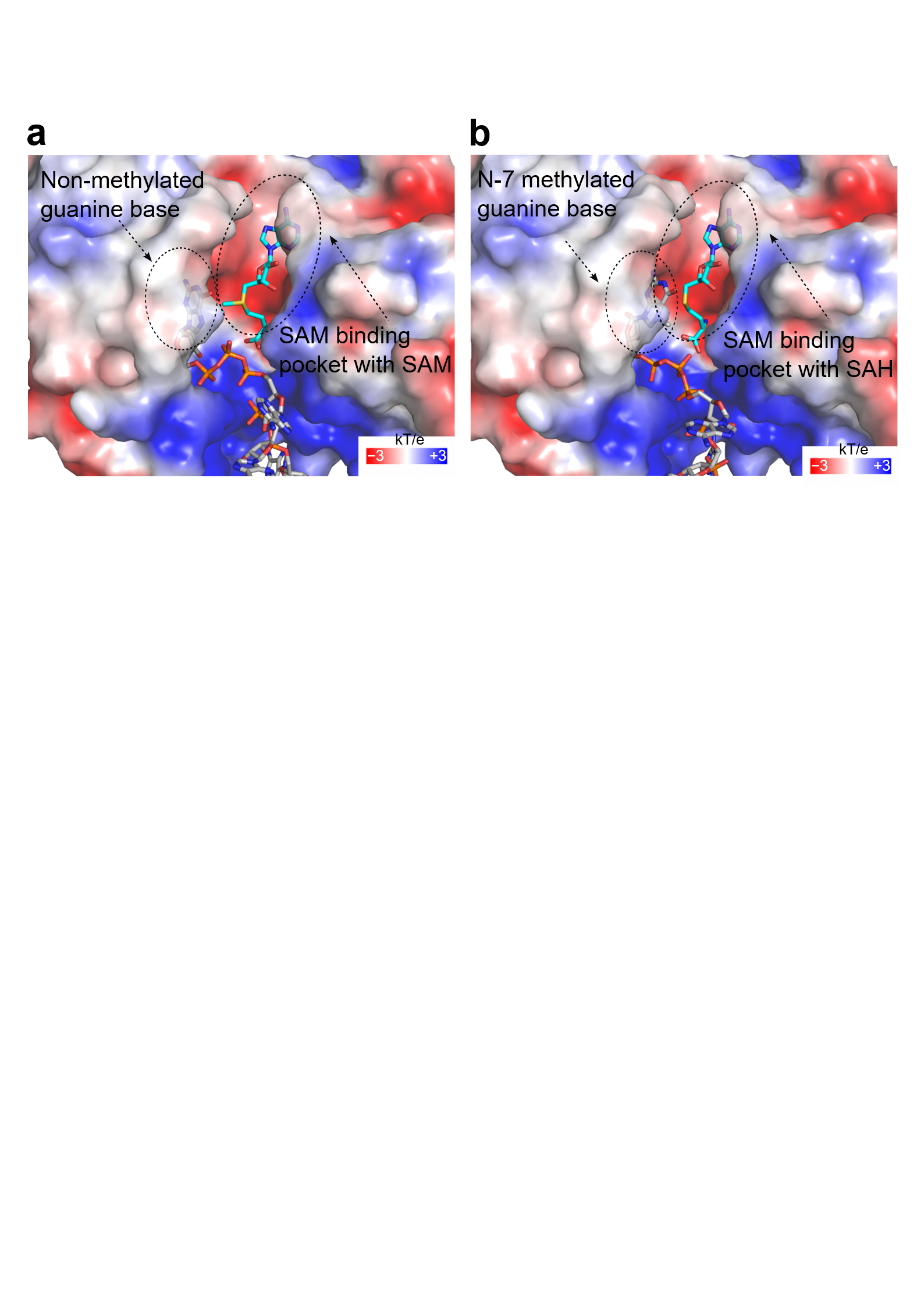

SI Figure 3 - **Model of non-methylated capped-RNA vs N7 methylated capped-RNA bound to the TvRNMT1**, semitransparent surface of the *Tv*RNMT1 protein is coloured according to the electrostatic surface potential: negative electrostatic potential is assigned to the red colour, while a positive electrostatic potential is assigned to the blue colour. The GpppG-capped RNA and m7GpppG-capped RNAs were modelled by AlphaFold/Boltz [1]. The coordinates of SAH molecule in the SAM binding pocket were used as a template (PDB:30VJ) for the precise SAM molecule position. SAH and SAM molecules are in stick representation and coloured in cyan. GpppG-capped RNA and m7GpppG-capped RNAs are shown in stick representation in silver colour and coloured according to the elements.

Table S1 - **X-ray data collection and refinement statistics**

| **Crystal name** | ***Tv*RNMT1**  **SAH** | ***Tv*RNMT1**  **sinefungin** | ***Tv*RNMT1**  **OBO101** |
| --- | --- | --- | --- |
| wavelength (Å) | 1.0596 | 1.0596 | 1.0596 |
| temperature (K) | 100 | 100 | 100 |
| space group | *P 1 2 1* | *P 1 2 1* | *P 1 2 1* |
| *a, b, c* (Å) | 53.78, 39.31, 85.35 | 54.00, 38.89, 85.06 | 54.04, 39.46, 84.75 |
| *α, β, γ* (deg) | 90.00, 90.92, 90.00 | 90.00, 90.77, 90.00 | 90.00, 90.48, 90.00 |
| resolution (Å) *^a^* | 45.83 - 1.60  (1.66 - 1.60) | 45.87 - 2.10 (2.176 - 2.10) | 45.74 - 1.89 (1.958 - 1.89) |
| number of unique reflections | 43 056 (3168) | 20 752 (1948) | 29 010 (2856) |
| multiplicity | 6.5 (4.6) | 6.5 (6.8) | 6.4 (6.3) |
| completeness (%) | 91.85 (68.22) | 98.64 (92.89) | 99.91 (100) |
| R_merge_*^b^* | 0.078 (1.426) | 0.221 (0.747) | 0.153 (1.057) |
| mean I/σ (I) | 11.67 (0.79) | 7.63 (2.55) | 8.61 (1.61) |
| CC_1/2_*^c^* (%) | 99.8 (40.3) | 98.9 (61.5) | 99.6 (75.5) |
| Wilson B (Å^2^) | 23.37 | 21.91 | 25.41 |
| **Refinement Statistics** | | | |
| number of reflections in  working set | 40 871 (3167) | 20 738 (1948) | 29 009 (2856) |
| number of reflections in test  set | 2185 (157) | 1038 (98) | 1450 (143) |
| *R* value*^d^* (%) | 19.21 (36.14) | 24.86 (33.70) | 22.91 (35.19) |
| *R*_free_ value*^e^* (%) | 22.34 (38.64) | 26.14 (33.30) | 25.85 (38.23) |
| number of molecules in AU*^f^* | 1 | 1 | 1 |
| number of atoms in AU*^g^*  protein/ligand/solvent | 2531 /46/158 | 2408/51/168 | 2525/56/136 |
| average ADP*^g^* for  protein/ligand/solvent (Å^2^) | 26.15 /22.1/33.32 | 26.18 /20.0/29.14 | 33.40/25.4/36.51 |
| RMSD bond length (Å) | 0.006 | 0.011 | 0.004 |
| RMSD bond angle (deg) | 0.82 | 1.30 | 0.61 |
| Ramachandran plot statistics*^h^* |  |  |  |
| favoured regions (%) | 98.41 | 97.67 | 97.13 |
| allowed regions (%) | 1.27 | 1.66 | 2.55 |
| **PDB code** | **30VJ** | **30VK** | **30VL** |

*^a^*Numbers in parentheses refer to the highest-resolution shell. *^b^R*_merge_=100∑*_hkl_*∑*_i_|*I*_i_*(*hkl*) – ‹I(*hkl*)›|/∑*_hkl_∑_i_*I*_i_*(*hkl*), where I*_i_*(*hkl*) is an individual intensity of the *i*^th^ observation of reflection *hkl* and ‹I(*hkl*)› is the average intensity of reflection *hkl* with summation over all data. *^c^*CC_1/2_ is the percentage of correlation between intensities from random half-datasets. *^d^*R-value = ||*F_o_*| – |*F_c_*||/|*F_o_*|, where *F_o_* and *F_c_* are the observed and calculated structure factors, respectively. *^e^*R_free_ is equivalent to R value but is calculated for up to 5% of the reflections chosen at random and omitted from the refinement process. *^f^*AU, asymmetric unit. *^g^*ADP, atomic displacement parameter, formally B-factor. *^h^*As determined by Molprobity [2].

**Supplementary Materials and Methods**

*Organic chemistry* - OBO101 was syntheiszed according to Scheme 1:
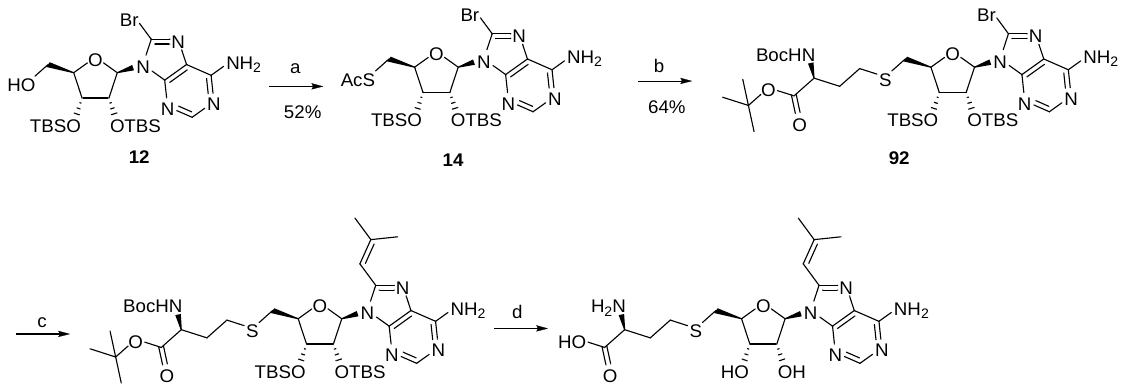

**Scheme 1.** Reagents, conditions, and yields: (a) PPh_3_, DIAD, AcSH, THF, 0 °C to rt; (b) “alkylbromide”, DBU, MeOH, 0 °C to rt; (c) 2-methyl-1-propenylboronic acid pinacol ester, Cs_2_CO_3,_ Pd(dppf)Cl_2_·DCM, dioxane, water, 90 °C, 52%; (d) TFA, water.

**((2*R*,3*R*,4*R*,5*R*)-5-(6-Amino-8-bromo-9*H*-purin-9-yl)-3,4-bis((tert-butyldimethylsilyl)oxy)tetrahydrofuran-2-yl)methanol (12)** was obtained as described previously (WO2006113615 A2 2006-10-26).

*S*-(((2*S*,3*R*,4*R*,5*R*)-5-(6-Amino-8-bromo-9*H*-purin-9-yl)-3,4-bis((tert-butyldimethylsilyl)oxy)tetrahydrofuran-2-yl)methyl) ethanethioate (14)

DIAD (2.10 mL, 10.7 mmol) was added dropwise over 5 min to an ice-cold solution of triphenylphosphine (2.83 g, 10.8 mmol) in dry THF (24 mL). After stirring for 30 min in the ice bath adenosine derivative **12** (3.00 g, 5.22 mmol) solution in THF (8 mL) was added, and the stirring was continued for 10 min at 0 °C. A solution of thioacetic acid (0.75 mL 10.5 mmol) in dry THF (2 mL) was added dropwise to the resulting yellow suspension and stirring was continued for another 1 h at 0 °C, then allowed to warm to room temperature and stirred for 3 h. The solvent was removed under reduced pressure. The residue was purified by chromatography on silica gel, eluent EtOAc in cyclohexane, gradient 1:3 - 1:2, chromatography on silica gel, eluent acetone in DCM 0-15% to obtain 1.71 g (52%) of the title compound as a beige solid.

^1^H NMR (400 MHz, CDCl_3_) δ 8.29 (s, 1H), 6.39 (s, 2H), 6.02 (d, *J* = 6.8 Hz, 1H), 5.46 (dd, *J* = 6.8, 4.2 Hz, 1H), 4.25 (dd, *J* = 4.2, 1.4 Hz, 2H), 4.14 – 4.06 (m, 1H), 3.56 (dd, *J* = 13.9, 6.9 Hz, 1H), 3.25 (dd, *J* = 13.9, 7.6 Hz, 1H), 2.34 (s, 3H), 0.96 (s, 9H), 0.78 (s, 9H), 0.15 (s, 3H), 0.14 (s, 3H), -0.07 (s, 3H), -0.41 (s, 3H).

^13^C NMR (100 MHz, CDCl_3_) δ 195.2, 153.8, 151.5, 150.8, 129.4, 120.7, 90.8, 85.3, 75.0, 72.0, 31.5, 30.6, 26.0 (3C), 25.8 (3C), 18.2, 17.9, -4.4 (2C), -4.5, -5.2.

HRMS (ESI^+^) *m/z* calcd for [M+H]^+^ C_24_H_43_O_4_N_5_BrSSi_2_ 632.1752, found 632.1750.

*tert*-Butyl *S*-(((2*S*,3*R*,4*R*,5*R*)-5-(6-amino-8-bromo-9*H*-purin-9-yl)-3,4-bis((*tert*-butyldimethylsilyl)oxy)tetrahydrofuran-2-yl)methyl)-*N*-(*tert*-butoxycarbonyl)-*L*-homocysteinate (92)

*tert*-Butyl (*S*)-4-bromo-2-((*tert*-butoxycarbonyl)amino)butanoate (0.33 g, 0.99 mmol) and nucleoside thioacetate derivative **14** (0.60 g, 0.87 mmol) were dissolved in dry MeOH (6 mL) under argon and the resulting suspension was cooled in the ice bath. DBU (0.19 mL, 1.27 mmol) was added, and the reaction mixture was stirred for 1 h at 0 °C, then allowed to warm to room temperature and stirred for 4 h. The reaction mixture was quenched by sat. NH_4_Cl, extracted with EtOAc, washed with brine, dried over Na_2_SO_4_ and concentrated. The residue was purified by chromatography on silica gel, eluent EtOAc in cyclohexane 20-100% to obtain 0.53 g (calculated yield 68%) of the title product as white foam. Contains 0.5 eq of diisopropyl hydrazine-1,2-dicarboxylate (11%) as an impurity.

^1^H NMR (400 MHz, CDCl_3_) δ 8.28 (s, 1H, H2), 5.98 (d, *J* = 6.1 Hz, 1H, H1′), 5.79 (br s, 2H, N^6^H_2_), 5.46 (dd, *J* = 6.1, 4.3 Hz, 1H, H2′), 5.05 (d, *J* = 7.7 Hz, 1H, NH), 4.44 (dd, *J* = 4.3, 2.4 Hz, 1H, H3′), 4.27 – 4.20 (m, 1H, Hα), 4.13 (td, *J* = 7.3, 2.4 Hz, 1H, H4′), 3.06 (dd, *J* = 13.9, 7.6 Hz, 1H, Ha5′), 2.88 (dd, *J* = 13.9, 6.6 Hz, 1H, Hb5′), 2.63 – 2.47 (m, 2H, Hγ), 2.10 – 1.97 (m, 1H, Haβ), 1.90 – 1.73 (m, 1H, Hbβ), 1.43 (s, 9H, C(CH_3_)_3_), 1.43 (s, 9H, C(CH_3_)_3_), 0.96 (s, 9H, Si-C(CH_3_)_3_), 0.79 (s, 9H, Si-C(CH_3_)_3_), 0.17 (s, 3H, Si-CH_3_), 0.15 (s, 3H, Si-CH_3_), -0.08 (s, 3H, Si-CH_3_), -0.38 (s, 3H, Si-CH_3_).

^13^C NMR (101 MHz, CDCl_3_) δ 171.4 (COO*t*Bu), 155.5 (COO*t*Bu^Boc^), 154.1 (C6), 152.2 (C2), 150.9 (C4), 129.2 (C7), 120.7 (C5), 91.0 (C1′), 85.4 (C4′), 82.3, 79.9 (2 x C(CH_3_)_3_), 74.8(C3′), 72.0 (C2′), 53.5 (Cα), 34.3 (C5′), 33.3 (Cβ), 28.6 (Cγ), 28.1, 27.1, 26.0, 25.8 (4 x C(CH_3_)_3_), 18.2, 18.0 (2 x Si-C(CH_3_)_3_), -4.3, -4.3, -4.4, -5.1 (4 x Si-CH_3_).

HRMS (ESI^+^) *m/z* calcd for [M+H]^+^ C_35_H_64_O_7_N_6_BrSSi_2_ 847.3274, found 847.3278.

*tert*-Butyl *S*-(((2*S*,3*R*,4*R*,5*R*)-5-(6-amino-8-(2-methylprop-1-en-1-yl)-9*H*-purin-9-yl)-3,4-bis((*tert*-butyldimethylsilyl)oxy)tetrahydrofuran-2-yl)methyl)-*N*-(*tert*-butoxycarbonyl)-*L*-homocysteinate (100)

8-Bromoadenosine derivative **92** (0.25 g, 0.30 mmol), Cs_2_CO_3_ (0.29 g, 0.88 mmol) were combined in a vial under argon, then degassed dioxane-water mixture (3:1, 3 mL) was added followed by Pd(dppf)Cl_2_·DCM (15 mg, 0.018 mmol, 6mol%) and 2-methyl-1-propenylboronic acid pinacol ester (0.10 mL, 0.49 mmol). The vial was sealed and the mixture was stirred for 18 h at 90 °C. The reaction mixture was diluted with EtOAc, washed with sat. NH_4_Cl, brine, dried over anh. Na_2_SO_4_ and concentrated. The residue was purified by chromatography on silica gel, eluent EtOAc in cyclohexane 15-100% to obtain 0.13 g (52%) of the title product as white foam.

^1^H NMR (400 MHz, CDCl_3_) δ 8.26 (s, 1H, H2), 6.22 – 6.19 (m, 1H, H1″), 5.85 (d, *J* = 5.8 Hz, 1H, H1′), 5.74 (s, 2H, NH_2_), 5.37 (dd, *J* = 5.8, 4.6 Hz, 1H, H2′), 5.05 (d, *J* = 8.1 Hz, 1H, NH), 4.48 – 4.43 (m, 1H, H3′), 4.28 – 4.17 (m, 1H, Hα), 4.11 (td, *J* = 7.0, 2.8 Hz, 1H, H4′), 3.07 (dd, *J* = 13.8, 7.4 Hz, 1H, Ha5′), 2.87 (dd, *J* = 13.9, 6.6 Hz, 1H, Hb5′), 2.62 – 2.50 (m, 2H, Hγ), 2.11 (d, *J* = 1.2 Hz, 3H, CH_3_3″), 2.06 – 1.98 (m, 1H, Haβ), 2.01 (d, *J* = 1.2 Hz, 3H, CH_3_3″), 1.88 – 1.76 (m, 1H, Hbβ), 1.42 (s, 9H, C(CH_3_)_3_), 1.23 (s, 9H, C(CH_3_)_3_), 0.95 (s, 9H, Si-C(CH_3_)_3_), 0.75 (s, 9H, Si-C(CH_3_)_3_), 0.17 (s, 3H, Si-CH_3_), 0.15 (s, 3H, Si-CH_3_), -0.10 (s, 3H, Si-CH_3_), -0.42 (s, 3H, Si-CH_3_).

^13^C NMR (101 MHz, CDCl_3_) δ 171.4 (COO*t*Bu), 155.5 (COO*t*Bu^Boc^), 154.5 (C6), 151.5 (C2), 150.1, 150.1 (C4 and C8), 149.4 (C2″), 119.9 (C5), 111.4 (C1″), 88.9 (C1′), 84.7 (C2′), 82.3 (C3′), 75.2, 74.9 (2 x C(CH_3_)_3_), 72.3 (C4′), 53.5 (Cα), 34.5 (C5′), 33.3 (Cβ), 28.7 (Cγ), 28.4, 28.1 (2 x C(CH_3_)_3_), 27.1 (C3″), 25.8, 25.0 (2 x C(CH_3_)_3_), 20.9 (C3″), 18.2, 18.0 (2 x Si-C(CH_3_)_3_), -4.2, -4.4, -5.2 (4 x Si-CH_3_).

HRMS (ESI^+^) *m/z* calcd for [M+H]^+^ C_39_H_71_O_7_N_6_SSi_2_ 823.4638, found 823.4645.

***S*-(((2S,3*S*,4*R*,5*R*)-5-(6-Amino-8-(2-methylprop-1-**en-1-yl)-9*H*-purin-9-yl)-3,4-dihydroxytetrahydrofuran-2-yl)methyl)-*L*-homocysteine (101)

Protected nucleoside **100** (0.12 g, 0.15 mmol) was dissolved in TFA - water mixture (3:1, 0.8 mL) at 0 °C, allowed to warm to room temperature and stirred for 24 h. The reaction mixture was evaporated, dissolved in water (2 mL), neutralized with NH_4_OH to pH 8, then evaporated, co-evaporated with EtOH. The residue was purified on silica gel, eluent water in MeCN 10-50%, then freeze-dried to obtain 20 mg (31%) of the title compound as a white foam.

^1^H NMR (400 MHz, DMSO) δ 8.11 (s, 1H, H2), 7.56 (br s, 3H, NH_3_), 7.13 (s, 2H, N^6^H_2_), 6.35 – 6.30 (m, 1H, H1″), 5.83 (d, *J* = 5.9 Hz, 1H, H1′), 5.38 (br s, 2H, OH2′ and OH3′), 5.10 – 5.00 (m, 1H, H2′), 4.24 – 4.15 (m, 1H, H3′), 4.01 – 3.92 (m, 1H, H4′), 3.24 – 3.17 (m, 1H, Hα), 2.93 (dd, *J* = 13.8, 6.1 Hz, 1H, Ha5′), 2.79 (dd, *J* = 13.6, 6.8 Hz, 1H, Hb5′), 2.61 (t, *J* = 7.7 Hz, 2H, Hγ), 2.15 (d, *J* = 0.9 Hz, 3H, CH_3_3″), 2.00 (d, *J* = 0.9 Hz, 3H, CH_3_3″), 1.99 – 1.92 (m, 1H, Haβ), 1.81 – 1.71 (m, 1H, Hbβ).

^13^C NMR (101 MHz, DMSO) δ 155.4 (C6), 152.0 (C2), 149.5 (C4), 147.9 (C8), 146.9 (C2″), 118.7 (C5), 111.7 (C1″), 88.1 (C1′), 83.3 (C4′), 72.5 (C3′), 71.0 (C2′), 53.1 (Cα), 33.8 (C5′), 31.4 (Cγ), 28.2 (Cβ), 26.7 (C3″), 20.6 (C3″).

HRMS (ESI^+^) *m/z* calcd for [M+H]^+^ C_18_H_27_N_6_O_5_S 439.1758, found 439.1758.

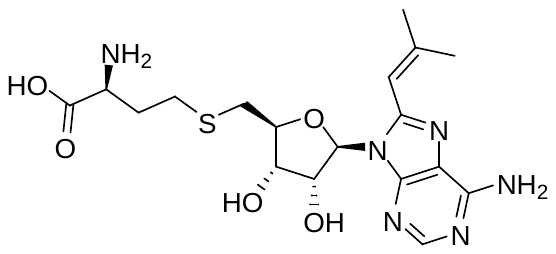

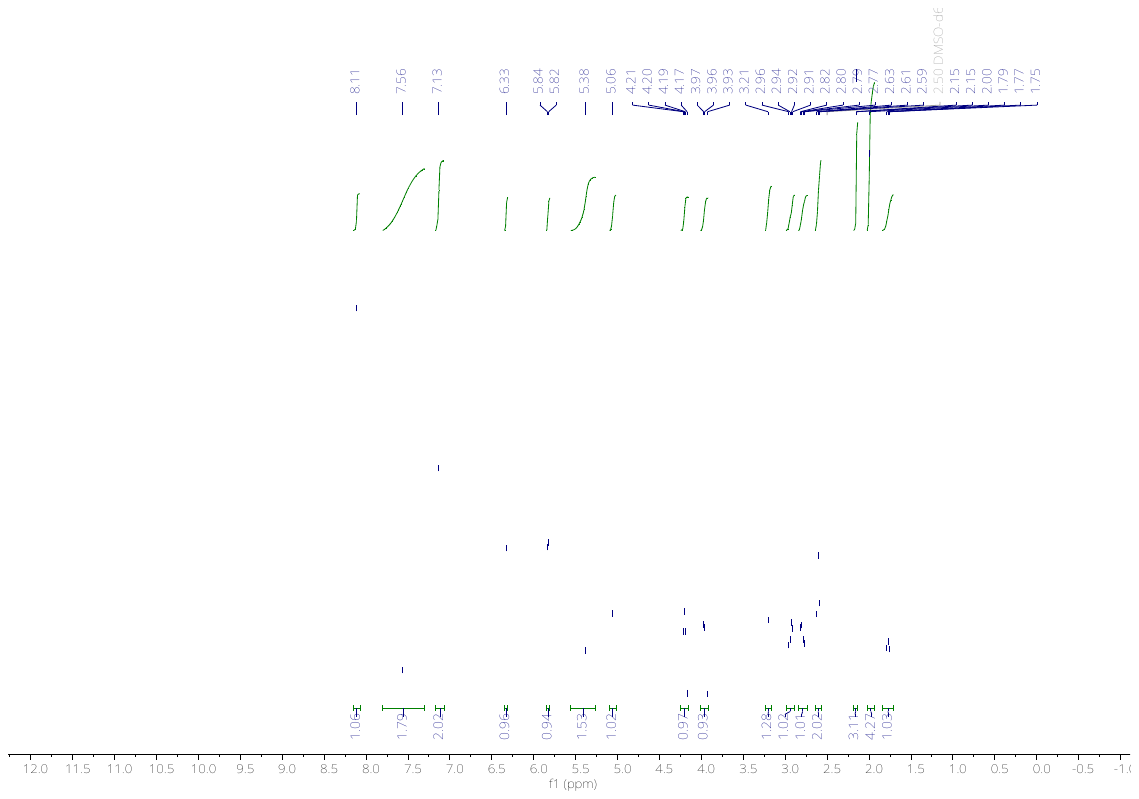

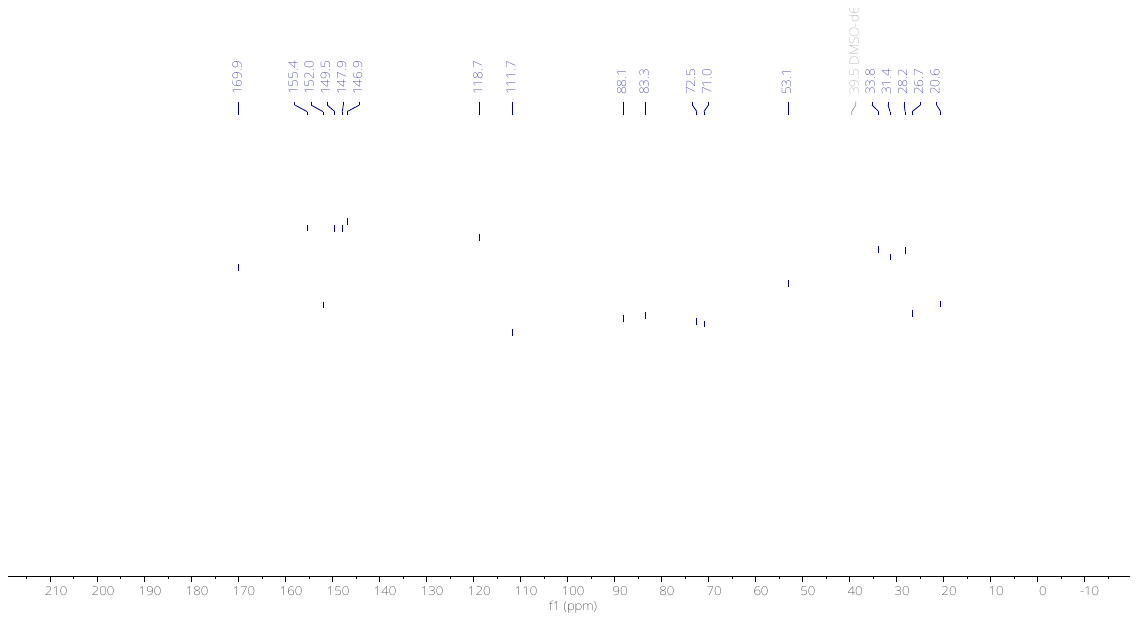

Graph 1 **- NMR spectra of OBO101**

Supplementary citations

1.	Wohlwend, J., G. Corso, S. Passaro, N. Getz, M. Reveiz, K. Leidal, W. Swiderski, L. Atkinson, T. Portnoi, and I. Chinn, *Boltz-1 democratizing biomolecular interaction modeling.* BioRxiv, 2025: p. 2024.11. 19.624167.

2.	Williams, C.J., J.J. Headd, N.W. Moriarty, M.G. Prisant, L.L. Videau, L.N. Deis, V. Verma, D.A. Keedy, B.J. Hintze, V.B. Chen, S. Jain, S.M. Lewis, W.B. Arendall, J. Snoeyink, P.D. Adams, S.C. Lovell, J.S. Richardson, and D.C. Richardson, *MolProbity: More and better reference data for improved all-atom structure validation.* Protein Science, 2018. **27**(1): p. 293-315.
